# Cytoscape stringApp: Network analysis and visualization of proteomics data

**DOI:** 10.1101/438192

**Authors:** Nadezhda T. Doncheva, John H. Morris, Jan Gorodkin, Lars J. Jensen

## Abstract

Protein networks have become a popular tool for analyzing and visualizing the often long lists of proteins or genes obtained from proteomics and other high-throughput technologies. One of the most popular sources of such networks is the STRING database, which provides protein networks for more than 2000 organisms, including both physical interactions from experimental data and functional associations from curated pathways, automatic text mining, and prediction methods. However, its web interface is mainly intended for inspection of small networks and their underlying evidence. The Cytoscape software, on the other hand, is much better suited for working with large networks and offers greater flexibility in terms of network analysis, import and visualization of additional data. To include both resources in the same workflow, we created stringApp, a Cytoscape app that makes it easy to import STRING networks into Cytoscape, retains the appearance and many of the features of STRING, and integrates data from associated databases. Here, we introduce many of the stringApp features and show how they can be used to carry out complex network analysis and visualization tasks on a typical proteomics dataset, all through the Cytoscape user interface. stringApp is freely available from the Cytoscape app store: http://apps.cytoscape.org/apps/stringapp.

## Introduction

Modern high-throughput technologies, including proteomics, produce an ever growing flow of new data on individual genes and proteins, which need to be interpreted in light of cellular context and existing biological knowledge. Protein network resources, in particular the STRING database^1^, have proven highly useful for providing such context. Indeed, such networks are very frequently shown in proteomics publications.

The STRING database provides known and predicted protein–protein associations data for a large number of organisms, including both physical interactions and functional associations with confidence scores that quantify their reliability. In addition to integrating available experimental data and pathways from curated databases, STRING predicts interactions based on co-expression analysis, evolutionary signals across genomes, automatic text-mining of the biomedical literature, and orthology-based transfer of evidence across organisms. However, the STRING web interface is not intended for large networks and provides limited flexibility in terms of network analysis and visualization, and accessing it without using the graphical user interface requires familiarity with programming.

The Cytoscape software^2,3^, on the other hand, is designed to analyze and visualize very large networks and provides much greater flexibility in terms of import of additional data and visualization of these onto networks. Moreover, Cytoscape has hundreds of apps, which users can install to add further functionality, such as clusterMaker2^4^ that implements numerous clustering algorithms and PTMOracle^5^ that allows PTMs to be analyzed in the context of protein networks. However, Cytoscape is a general network tool, not a network database, and as such needs to import its networks from elsewhere.

Together this makes STRING and Cytoscape a perfect match, especially for analysis of proteomics data; indeed, more than thousand papers in PubMed Central mention both STRING and Cytoscape, clearly demonstrating a strong need to integrate them into a single workflow. We have done exactly that by developing the stringApp, a Cytoscape app that facilitates import of STRING networks into Cytoscape and integration with additional user-provided data. At the same time, the app provides the look and many of the features of the STRING web interface within Cytoscape. The app supports several types of queries to retrieve networks starting from either a list of proteins, a disease of interest from the DISEASES database^6^, or a PubMed query. Moreover, it provides access to additional data from associated resources, namely small molecule interactions from STITCH^7^, subcellular localization from COMPARTMENTS^8^, tissue expression from TISSUES^9^, and drug target information from Pharos^10^. Together, these features enable users to easily carry out complex network analysis and visualization tasks, all through the graphical user interface of Cytoscape. In a typical use case, we demonstrate how a proteomics dataset can be analyzed and visualized with the help of the stringApp and Cytoscape.

## Methods

### Data sources used by stringApp

The stringApp retrieves information collected from several source databases. The protein network is imported from the current STRING v10.5^1^ and augmented with protein–chemical and chemical–chemical associations from the current STITCH version 5^7^. This is complemented by drug-target classification from the current release of Pharos^10^ and information on disease associations, tissue expression, and subcellular localization from the weekly updated databases DISEASES^6^, TISSUES^9^, and COMPARTMENTS^8^.

Although these databases all provide Application Programming Interfaces (APIs), we mirror the data from the current production versions of STRING and STITCH in a dedicated PostgreSQL database on the same server that already hosts DISEASES, TISSUES, and COMPARTMENTS. This is done both to provide additional functionality over the existing APIs and to allow stringApp to efficiently retrieve all information for a protein network as with a single API request.

### Algorithms implemented at the database level

Another major benefit of having all data available in a single database is, that it allows us to implement certain algorithms, as described below, at the database level. Instead of first loading large amounts of data from one or more databases into memory and then executing the algorithms, we were able to implement the algorithms in Structured Query Language (SQL) and execute them directly within the PostgreSQL database engine. We made use of this approach for two algorithms used by stringApp.

#### Network expansion

This algorithm adds *N* additional nodes to the network based on their total connectivity to a current selection of nodes (*X*) relative to their overall connectivity in the STRING database (if no nodes are selected, the complete network is considered as the selection). All nodes not currently in the network are ranked according to the following score:

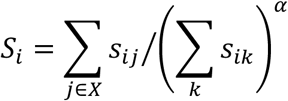

where *S_ij_* is the confidence score between the node *i* (to be considered for inclusion in the network) and the node *j* (in the current selection of nodes (X)), while the sum over *s*_*ik*_ captures the connectivity of node *i* to all other nodes *k* in the database. The parameter *α* is called the *selectivity* in the stringApp user interface (Expand network option) and has a default value of *0.5*. This value gives a good tradeoff between choosing nodes that have high confidence links to the selection but possibly also to many other proteins (low selectivity), and choosing nodes that are specifically linked to the selected nodes but with lower confidence (high selectivity). The sum in the enumerator is calculated on-the-fly using SUM aggregate function in SQL, whereas the sum in the denominator has been precalculated for all nodes in the aforementioned database. We can thus with a single SQL command score all candidate nodes, rank them, and return the top *N*.

#### PubMed query

The second algorithm implemented in SQL is used to retrieve a network based on a PubMed query. The app sends the user-specified query to the PubMed API to retrieve the set (*X*) of matching PMIDs, selects the top *N* entities that are preferentially mentioned in *X*, and finally retrieves the network for them. To rank the entities, we use the following scoring function:

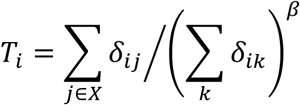

where *δ_ij_* is 1 if the molecular entity *i* is mentioned in abstract *j* and 0 otherwise, while *δ_ik_* is 1 if the molecular entity *i* is mentioned in any abstract *k* in PubMed and 0 otherwise. The parameter *β* is fixed to a value of 0.4 based on previously published text-mining experiments^6^. It serves a similar purpose to the selectivity described above, controlling the tradeoff between choosing entities that are mentioned in as many of the selected abstracts as possible but possibly also in many other abstracts vs. choosing entities that are specifically mentioned only in the selected abstracts. Given the similarity of this formula to the one used for network expansion, it should come as no surprise that it too can be implemented as a single SQL command which calculates the enumerator using the COUNT aggregate function, whereas the denominator has been precalculated for all pairs of entity and abstracts in the aforementioned database.

### Implementation of the app

The stringApp is implemented in Java utilizing the Cytoscape 3.6 App API. The app has two main functions: (1) to serve as a bridge between Cytoscape and the web service APIs of STRING and the related databases, and (2) to provide visualizations resembling the ones on the STRING web server as well as additional features like the side panel and enrichment visualizations. These two functions work together to bring much of the richness of the STRING web site into Cytoscape, which then allows the network and all associated data to be analyzed with Cytoscape and its hundreds of other apps. For instance, the clusterMaker2 app^4^ can be very useful for clustering STRING networks, as shown in the use case below.

The bridge functionality of the stringApp uses several RESTful^11^ web service APIs to query the databases and retrieve networks. In case of protein and protein/compound queries, the app first resolves the entered query terms to the internal database identifiers using the standard STRING and STITCH API. For disease queries, it instead contacts the API of the DISEASES database twice, first to resolve the entered disease name to a disease identifier, and second to retrieve the list of proteins associated with the disease. For all three types of queries, stringApp provides the user with the ability to manually resolve any ambiguous names. The handling of PubMed queries was described in the previous section. Irrespective of the type of query, these steps result in a list of nodes, for which stringApp retrieves all node and edge data by calling the web service API of the dedicated PostgreSQL database. The latter API is also used to retrieve any node or edge data required when expanding an existing network, lowering the confidence cutoff, or adding additional nodes to a network.

The stringApp retrieves functional enrichment analysis results for a whole STRING network or a selected subset of it by sending a request to the STRING enrichment API. The results are stored and shown in a Cytoscape table called *STRING Enrichment*, which lists all enriched terms along with their gene counts, corresponding FDR values, and gene sets. Since the list of enriched terms can become very long, especially for large networks, the app allows the user to filter the enrichment results to show terms from any combination of six term categories as well as to eliminate redundant terms, which represent similar sets of genes.

The redundancy filtering takes the list of enriched terms sorted by FDR value and removes the terms that are too similar to any of the previous, better scoring terms that were not themselves removed (also referred to as the Hobohm 1 method^12^). The similarity between two terms is measured by the Jaccard index of the sets of genes annotated by the two terms. A term is added to the filtered list only if it has Jaccard similarity less than the user-specified redundancy cutoff to any other term already in the filtered list.

To retain the look and feel of STRING networks, the stringApp adds a new *STRING* Visual Style to the already existing set of Cytoscape styles. This style enables the glass ball effect and the optional visualization of the protein or compound structures within the nodes. These visual properties can be enabled or disabled by the user from the stringApp menu. The initial node colors are assigned arbitrarily by the app but can be easily substituted by a node color mapping of any node attribute. In addition to the node visual properties, the STRING style also includes a mapping of the interaction confidence scores to edge color and thickness.

### Specific data and software for the use case

The proteomics data used in the case study were obtained from a phosphoproteomics study of ovarian cancer^13^ (specifically Suppl. Table 3 of the study). The list of proteins used to retrieve the network was extracted from the significantly regulated phosphorylation sites listed in this table. Furthermore, log-ratios of abundance in disease versus healthy tissues were computed based on the average abundance values over the samples listed in the same table. To facilitate the subsequent visualization in Cytoscape, we also modified the Suppl. Table 3 by keeping only the significantly regulated phosphorylation sites and sorting them by significance. This modified version of the table, which was imported into Cytoscape, is provided as Supporting Information 2.

During import of the associated log-ratios and phosphorylation cluster assignments, the most significant phosphorylation site was chosen whenever multiple sites were found on the same protein (by first sorting the table based on “Gene name” and then on “adj. p-value”).

All analyses were performed on the 12th of April 2018 using Cytoscape version 3.6.1 and stringApp version 1.3.2 and are provided in a Cytoscape session (DOI: 10.6084/m9.figshare.7258235). Additionally, we used clusterMaker2 version 1.2.1 to perform Markov clustering (MCL)^14^ of the protein network and EnhancedGraphics version 1.2.0^15^ to enable stringApp visualization of enriched terms as circular plots onto the network nodes.

## Results

### Presentation of the stringApp

The stringApp was designed to serve as a bridge between two well-known and widely used resources, the STRING database for quality-controlled protein–protein association networks and the Cytoscape software platform for network data integration, analysis and visualization. Thus, the core purpose of the stringApp is to retrieve network data from STRING, import it into Cytoscape and retain the look and most of the functionality of the STRING database, while at the same time allowing users to analyze the network with the full set of Cytoscape features and integrate it with their own data. Nevertheless, stringApp also imports protein-protein interactions from STRING for a disease or PubMed query of interest as well as protein–chemical interaction data from STITCH. A list of the main stringApp features can be found in Table 1.

**Table 1.** Main stringApp features with corresponding inputs and outputs.

| **Function** | **Operation** | **Input** | **Output** |
| --- | --- | --- | --- |
| *STRING:*<br>*protein query* | Create a network for one or multiple proteins | Protein name(s) or identifier(s) | Network |
| *STRING:*<br>*disease query* | Create a network of proteins associated with a disease | Disease name or identifier |  |
| *STRING:*<br>*PubMed query* | Create a network by entering a PubMed query | PubMed query |  |
| *STITCH:*<br>*protein /*<br>*compound query* | Create a network for one or multiple proteins and compounds | Protein/compound name(s) or identifier(s) |  |
| *Expand network* | Expand network by more interactors | STRING network<br>Number and type of interactors to expand by | Network with nodes and edges added |
| *Query for additional nodes* | Add more proteins or compounds to an existing network | STRING network<br>Protein/compound name(s) or identifier(s) |  |
| *Change confidence* | Change confidence of the network | STRING network<br>New confidence cutoff | Network with edges added or removed |
| *Retrieve functional enrichment* | Retrieve functional enrichment for several categories | STRING network | Enrichment table |
| *Filter enrichment* | Filter terms in the enrichment table by category or redundancy | STRING enrichment table<br>Filter options | Filtered enrichment table |
| *Draw charts* | Show selected enrichment terms as charts on the nodes | STRING network<br>Enrichment table | Network with enrichment visualization |

Currently, four different types of queries are supported by the stringApp, which allow users to retrieve a STRING network starting from 1) a list of one or more genes/proteins, 2) a list of chemical compounds, 3) a disease, or 4) a PubMed query. Additionally, the user can choose the species of interest and the confidence cutoff for the interactions to be retrieved. The *STRING: protein query* obtains a STRING network for an arbitrarily long list of proteins and can be used, for example, to retrieve a STRING network for a proteomics or transcriptomics study. In a similar manner, the *STITCH: protein/compound query* obtains a network for a list of protein or chemical compound names from STITCH as shown in Fig. 1. The *STRING: disease query* first queries the DISEASES for the top-N human proteins associated with the disease specified by the user and then retrieves a STRING network for these. The *STRING: PubMed query* allows users to get a STRING network for any topic of interest by first querying PubMed for abstracts pertaining to the topic, then using text mining on these abstracts to identify the top-N proteins associated with the topic, and retrieving a STRING network for these.

**Figure 1.**
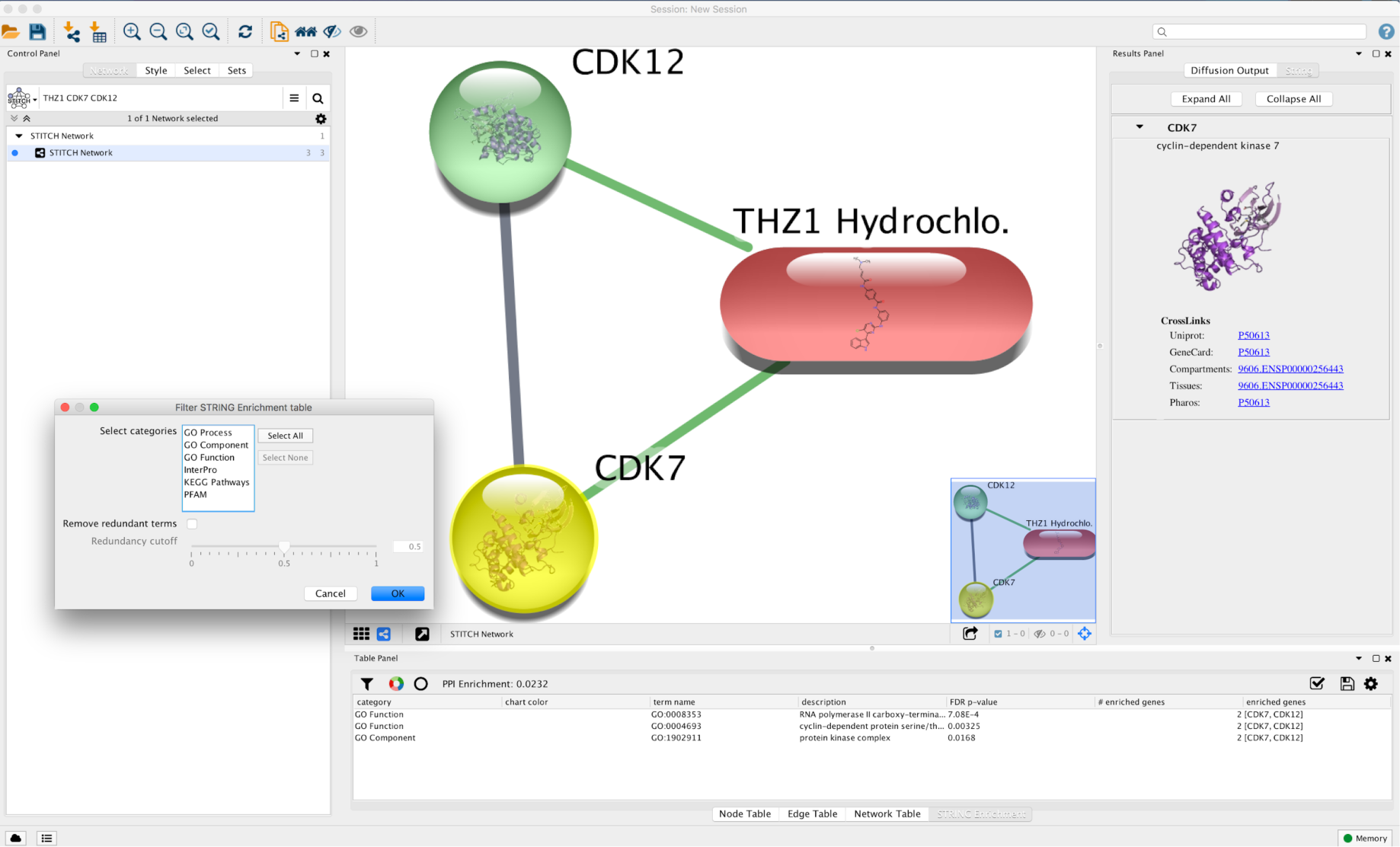
Highlighting various stringApp features in a screenshot of a small STITCH network in Cytoscape. Edge colors indicate type of interaction (green for protein-compound and grey for protein-protein interactions) and node colors are arbitrary. The results panel (right) shows the 3D structure of the currently selected node CDK7 (indicated by yellow node color) and provides links to other related resources. The STRING enrichment table panel (bottom) lists the enriched terms for this network (FDR-corrected p-value < 0.05) with their category, term name description, FDR-corrected p-value and the enriched genes. The Filter STRING Enrichment table dialog (left) demonstrates the available options for filtering enriched terms by category and redundancy.

In addition to the interactions from STRING/STITCH, stringApp retrieves a variety of related information, which is stored as node and edge attributes for each protein/chemical or interaction, respectively. The node attributes include the STRING and UniProt accession numbers, which allow for cross-linking with other resources, a human-readable name for display purposes, the protein sequence or a chemical SMILES string, and a structure image where possible. The edge attributes include the overall confidence score of each interaction as well as the subscores from each individual evidence channel in STRING/STITCH. Whenever available for the organism in question, information on the tissue expression and subcellular localization of each protein is included from the TISSUES and COMPARTMENTS databases. Furthermore, stringApp fetches drug target information from Pharos. If a protein was retrieved through a disease query or PubMed query, the corresponding confidence score for the disease–gene association according to DISEASES or text-mining score (see Methods) are included as node attributes. As shown in Fig. 1, stringApp also provides a results panel in Cytoscape, which shows the protein or compound structure of a selected node as well as links to other related resources, including UniProt^16^, GeneCards^17^, COMPARTMENTS, TISSUES, and Pharos.

Once a STRING/STITCH network is in Cytoscape, it can be modified in several ways. First, users can expand the network with the nodes that are most strongly connected with the nodes currently in the network or with a selected subset of them (see Methods for details on the underlying algorithm). These new nodes can be either proteins from STRING or chemical compounds from STITCH. Second, it is possible to add specific new nodes to the network by providing their names just like in the original query. Third, users can change the confidence cutoff of the imported interactions; increasing it filters the current network to remove edges that do not pass the new cutoff, whereas decreasing it will re-query the server to fetch the additional interactions, that did not pass the original cutoff.

Network analysis and functional enrichment analysis are complementary methods to gain an overview of a long gene or protein list. The stringApp allows users to combine the two, by first performing an enrichment analysis and subsequently visualizing the results onto a STRING network. To do so, the user specifies the enrichment significance threshold (with default value of 0.05). Then, enriched Gene Ontology terms, KEGG Pathways, and protein domains are retrieved from the STRING enrichment web service and shown as a table (see example in Fig. 1). From the table, the user can then optionally filter the enrichment results to reduce redundancy (see Methods for details) and visualize the top terms onto the network as donut or pie charts using ColorBrewer^18^ palettes to distinguish the different terms.

### Use case: Analysis of an MS-based phosphoproteomics dataset

To illustrate some of the more important features of stringApp that are relevant to analysis of proteomics data, we have chosen a typical dataset resulting from a phosphoproteomics study of ovarian cancer by Francavilla *et al.*^13^ published in 2017. In this study, the authors compare the phosphoproteome of primary cells derived from epithelial ovarian cancer (EOC) and two healthy tissues, namely ovarian surface epithelium (OSE) and distal fallopian tube epithelium (FTE). The aim of the study was to uncover cancer-specific changes in expression, phosphorylation state, and kinase signatures.

In the following sections, we will go through how this dataset can be analyzed and visualized in a variety of ways using the stringApp and Cytoscape. Starting with the list of proteins with significantly regulated phosphorylation sites in the study, we first retrieve the corresponding STRING network in Cytoscape. Then, we import data from the study, namely the log-ratios of phosphorylation between disease cells and healthy tissues and the phosphorylation cluster assignments, to be able to visualize them on the network nodes. To gain insight from the resulting highly connected protein network, we partition it using a clustering algorithm and relay out the network. The largest identified cluster turns out to be highly relevant to the main findings of the study since it contains both CDK7 and POLR2A as well as many splicing related proteins. The study by Francavilla *et al.* showed that CDK7 phosphorylates POLR2A and regulates EOC cell proliferation, and that peptides in proteins with splicing variants were overrepresented in the EOC proteome. We thus focus on this cluster, analyze it for enriched functional terms, and visualize selected terms on the network. Finally, we highlight the proteins from the study that are annotated as druggable targets in the Pharos database or associated with EOC according to the DISEASES database.

### Network retrieval and data import

The first step of the analysis is to retrieve a STRING network for the 541 unique proteins with significantly regulated phosphorylation sites. This is done by opening the *Import Network from Public Databases* dialog, choosing *STRING: protein query*, entering their UniProt accession numbers into the dialog, and leaving all query parameters at their default values. The resulting network retrieved from STRING 10.5 consists of 537 nodes and 3027 edges with the default confidence score of 0.4 and above, which is consistent with the default of the STRING web site. Four proteins in the dataset, three of which were unreviewed TrEMBL entries, could not be mapped to STRING identifiers by the app and were thus not included in the further analysis. To simplify the figures, we also opted to delete the 78 singleton nodes, i.e. the proteins with no interactions in the retrieved network.

To add data from the proteomics study to the STRING network, we import the modified version of Suppl. Table 3 from Francavilla *et al.* (Supporting Information 2) using the built-in functionality of Cytoscape to import data columns from a tabular file. In this step it is crucial to correctly choose which column from the file should be mapped to which column in the Cytoscape node table; if the identifiers do not match, the data will not be imported. To facilitate this mapping, stringApp saves the user-provided query identifiers in the “query term” column in the Cytoscape *Node table*. In this use case, the UniProt identifiers from the “Uniprot” column in the supplementary table (Supporting Information 2) were used to retrieve the STRING network and therefore, this is the column that should be selected as the “Key” column in the table preview in the *Table import* dialog. Since these identifiers were stored in the “query term” column in the Cytoscape *Node table*, it should be selected as the “Key Column for Network” in the *Import table* dialog. Upon successful import of the data, the new columns are inserted at the end of the Cytoscape *Node table*. Here, we import the average log-ratios between disease and healthy tissue (“EOC vs. EOS&FTE” column) and the phosphorylation cluster assignments (“Cluster” column).

### Network layout and visual mapping of data

Having imported the proteomics data, we can map it onto the nodes in the network using the Cytoscape Visual Styles functionality. Numeric data such as the log ratios between disease and healthy tissues are best shown using a continuous mapping of the values to a color gradient. Here, we use a blue–white–red color gradient to highlight nodes with low or high log ratios (Fig. 2). Categorical data such as the phosphorylation cluster assignments should be represented by a discrete color mapping, which assigns a different color to each category (Fig. 3). Mappings between visual properties and attributes can also be created for edges; the default STRING visual style uses this to show edges with higher confidence scores as thicker, darker lines.

**Figure 2.**
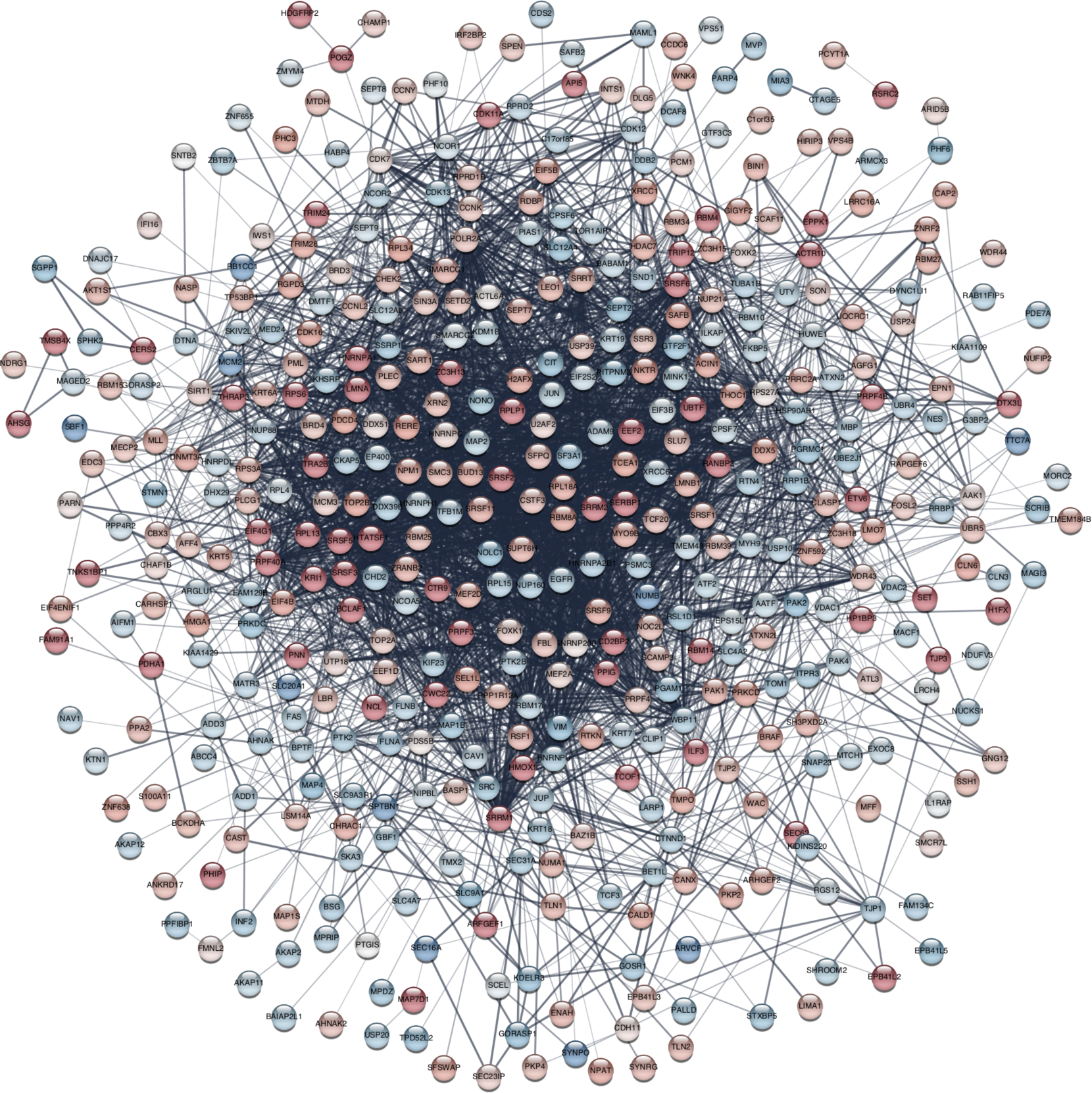
STRING network of proteins with significantly regulated phosphorylation sites detected in a phosphoproteomics study of ovarian cancer^13^. Log-ratios between disease and healthy tissues for the most significant site for each protein were mapped to the nodes using a blue-white-red gradient. Proteins without any interaction partners within the network (singletons) are omitted from the visualization.

**Figure 3.**
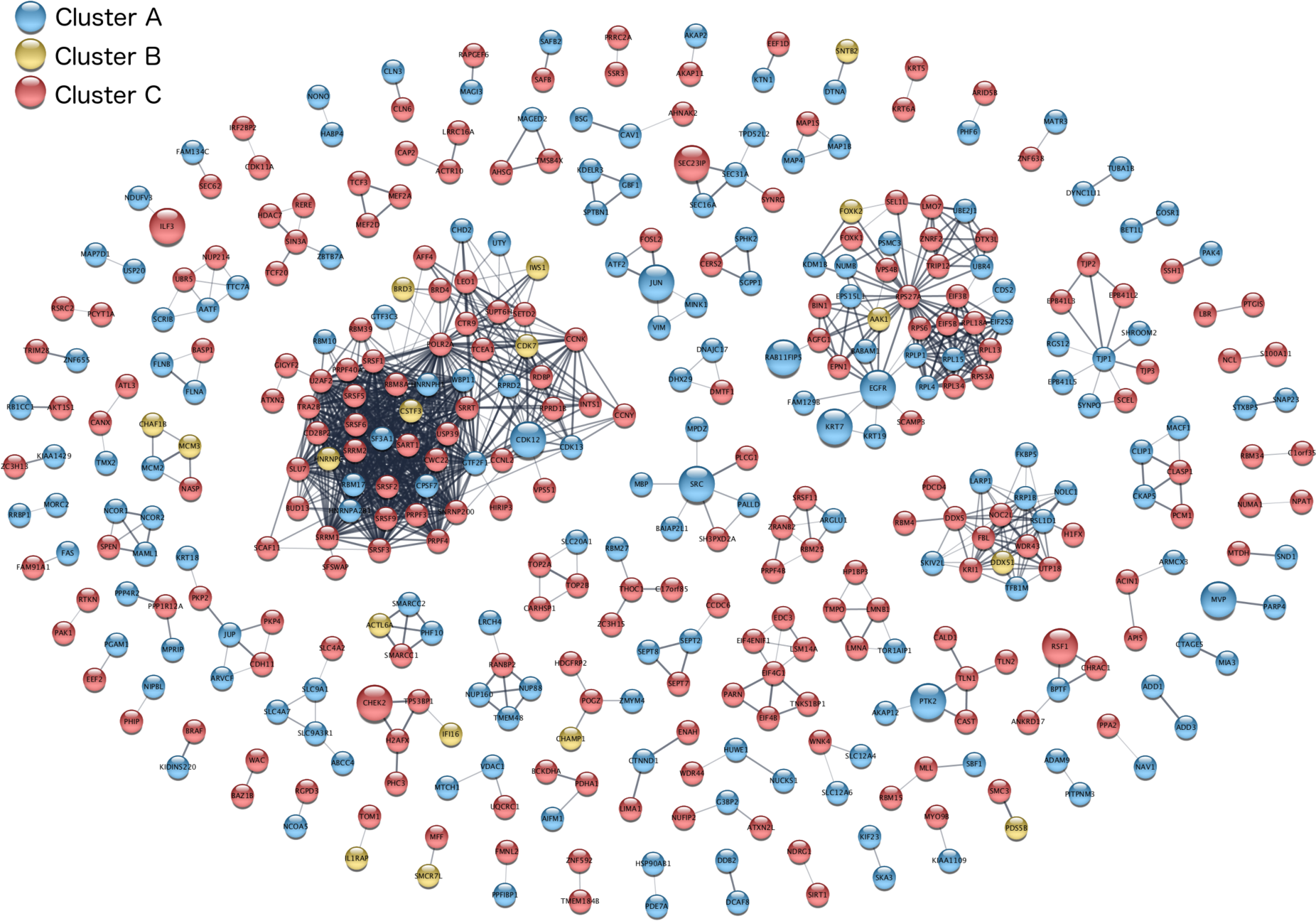
Clustered protein association network with proteins colored by the phosphorylation cluster to which they were assigned in the original analysis^13^. Clustering was performed using the Markov clustering (MCL) implementation in the clusterMaker2 Cytoscape app. The 13 proteins associated with *Ovary epithelial cancer* according to DISEASES are represented by bigger nodes. Clusters consisting of one node only are omitted from the visualization.

Network visualization of large proteomics datasets is challenging for several reasons. First, these networks tend to be large, typically consisting of hundreds to thousands of proteins with thousands to tens of thousands of interactions between them as exemplified by Fig. 2. Visualizing large, dense networks in a way that reveals the patterns within them, such as groups of similarly regulated proteins, is inherently difficult^19^. Second, whereas a single comparison of two conditions is easily visualized using a color gradient, many proteomics studies, including the one used in this example, compare multiple conditions or timepoints.

### Using clustering to improve visualization

Clustering can be a powerful strategy to visualize multidimensional data on large networks. In Cytoscape, a broad selection clustering algorithms are available through the widely used clusterMaker2 app^4^. This app can cluster the nodes in the network both based on the edges that connect them (network clustering) and based on numeric data from the Cytoscape *Node Table* (attribute clustering). We could thus have used the attribute-clustering algorithms in clusterMaker2 to identify groups of proteins that exhibit similar changes in phosphorylation. However, in this use case we instead opted to import the phosphorylation cluster assignments from the original study as described in the previous section.

To group the proteins in the network based on their interactions from STRING, we used clusterMaker2 to run Markov clustering (MCL)^14^. We increased the *inflation value* to 4.0 to reduce the cluster size, set *array sources* to use the STRING confidence *score* attribute as weights, checked the option to *create new clustered network*, and left all other settings at their default. The resulting network is greatly simplified and much easier to visualize, since only the 1058 interactions within clusters are retained (Fig. 3).

Finally, to visualize how the proteins are regulated, we color the nodes based on the which phosphorylation cluster they were assigned to: *cluster A* (blue) is up-regulated in both healthy tissues (FTE and OSE), *cluster B* (yellow) is up-regulated in one healthy tissue (FTE) and in disease tissue (EOC); and *cluster C* (red) is up-regulated in disease tissue (EOC). For comparison, we also provide the same network colored by log ratios between disease and healthy tissues (Supporting Information 1, Figure S1).

### Functional enrichment analysis

In the last parts of the use case we will focus on the largest cluster in the network, which consists of 62 proteins and relates several of the findings of the original study. We thus created a new, separate network in Cytoscape that consists only of this cluster.

To functionally characterize the cluster, we used stringApp to perform functional enrichment analysis with an FDR threshold of 5%, which resulted in a list of 129 statistically significant terms that span all six categories: GO Biological Process, GO Molecular Function, GO Cellular Component, KEGG Pathways, PFAM and InterPro protein domains. We next used the filter functionality to eliminate redundant terms (using the default redundancy cutoff of 0.5), thereby reducing the list to a more manageable 38 enriched terms. Of these, the two most significant terms were the GO biological process *mRNA processing* and the KEGG pathways *Spliceosome*, which covered 39 and 20 out of the 62 proteins in the cluster, respectively. To show which proteins are annotated with which of these terms, we used the stringApp to visualize them as a *split donut charts* around the nodes (Fig. 4.). These enrichment results fit well with the finding by Francavilla *et al.*^13^ that peptides from proteins with splicing variants were overrepresented among EOC-regulated phosphorylated peptides. Moreover, the same cluster contains the protein POLR2A, the phosphorylation of which has been associated with both transcriptional regulation and alternative splicing^20^.

**Figure 4.**
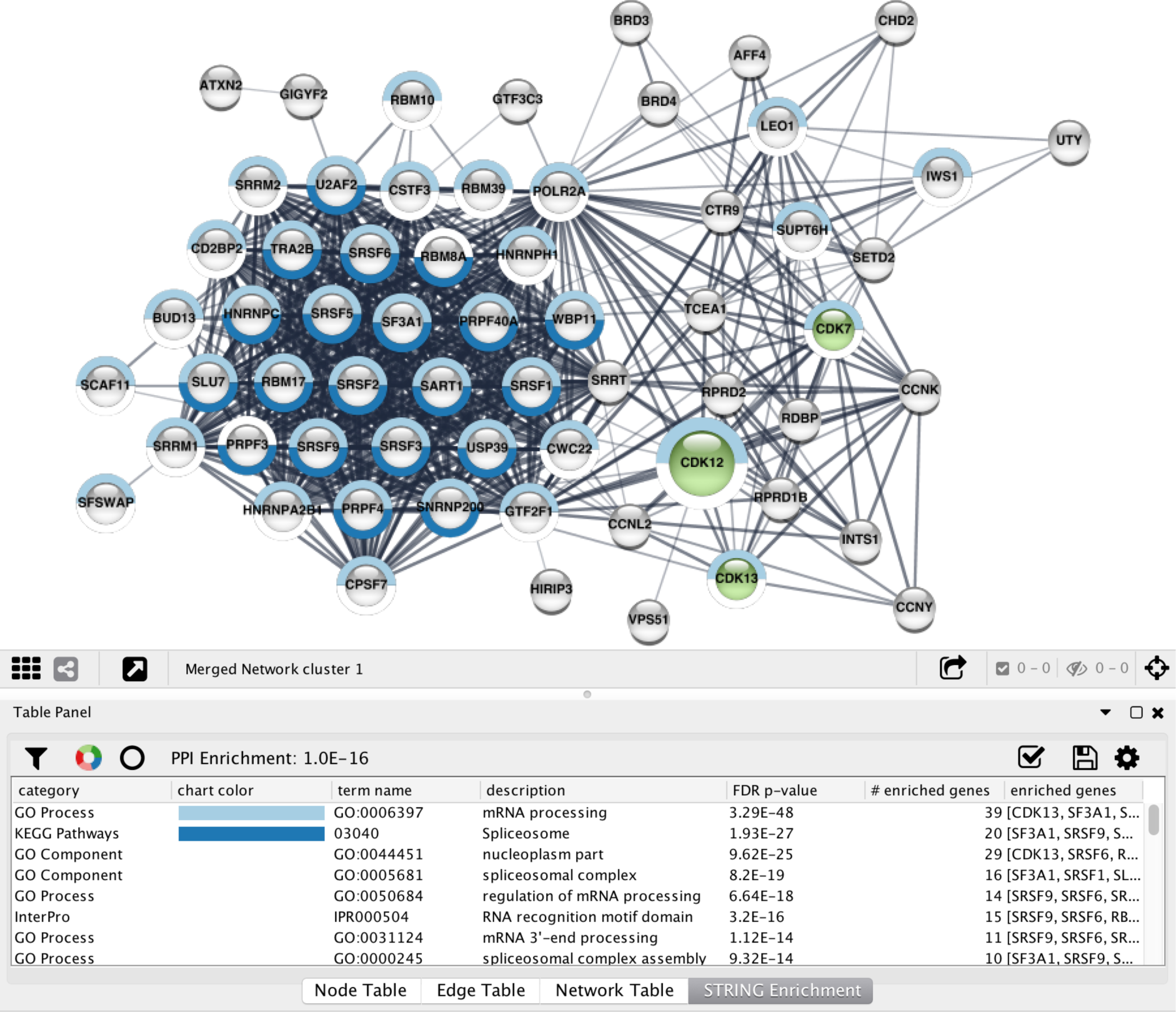
Functional analysis of the largest cluster obtained by Markov clustering (Fig. 3). The top-2 enriched terms after redundancy filtering were visualized as split donut charts around the nodes annotated with those terms. CDK12 is highlighted by bigger node size because it is associated with *Ovary epithelial cancer* according to DISEASES. The three kinases CDK7, CDK12 and CDK13 are highlighted in green based on annotations from the Pharos database.

### Annotation of disease-associated proteins and drug targets

The stringApp automatically retrieves drug target information from the Pharos database into the “target development level” and “target family” columns of the Cytoscape *Node table*. The latter column includes annotations of known kinases and other drug target families. Using the discrete mapping functionality of Cytoscape, one can highlight the kinases (and other drug target families) by assigning different colors to the corresponding nodes (see Fig. 4). The cluster contains 3 of the 22 kinases present in the full network, including CDK7 that the study showed phosphorylates POLR2A and thereby likely regulates the processes identified in the enrichment analysis.

Finally, to annotate the network with proteins already associated with EOC, we use the *STRING: disease query* functionality of stringApp to import a STRING network of the top 500 candidate disease genes according to the DISEASES database. The confidence scores of the associations between these genes and EOC range from 0.69 to 2.66 on a scale from 0 to 5. As a compromise between confidence and coverage, we decided to keep only genes with a disease confidence score above 1.0, resulting in a network of 222 genes likely associated with EOC. We then identified the proteins from the DISEASES network in the study network by first creating the union of the two networks using the *Merge Networks* tool in Cytoscape and then removed all nodes not coming from the study. In the resulting network, the node attribute *disease score* marks all proteins associated with EOC according to the DISEASES database, which we used to highlight them as bigger nodes (see Fig. 3 and 4).

## Discussion

### Scope of the stringApp

In our use case, we have illustrated how many of the features of stringApp (see Table 1 for a more comprehensive list) and Cytoscape can be used to analyze and visualize a proteomics study of human cells. However, this does not showcase the full scope of the app.

The current version of STRING provides functional association networks for more than 2000 different organisms, all of which can be accessed through stringApp. Moreover, the app can equally well be used to visualize other types of high-throughput experiments that give rise to a list of genes or proteins. This is true for transcriptomics data, which can be imported and visualized on a network by following the same steps we showed for proteomics data, as well as for phenotypic screens and mutation data. For example, the stringApp has already been used in the literature for network analysis of microarray data on the mammalian circadian pacemaker^21^ and for comparing a coexpression network obtained from maize RNA-seq data to a network from STRING^22^.

One key feature of the stringApp, which was not used in the use case, is the ability to expand a network. This uses the STRING network as a whole to bring in additional proteins that were not initially identified, but which may be of interest because they are preferentially associated with the proteins on the initial list. In case of phosphoproteomics data, this can be used to identify proteins that may not themselves be regulated through phosphorylation, but which function together with proteins that are.

While users can import their own data, it is worth noting the stringApp also automatically augments the network with tissue expression data and information on protein subcellular localization data on tissue expression. Without having to provide any data, it is thus possible for users to visualize which proteins localize to a certain tissue or part of the cell. The data can also be highly useful for filtering networks to produce, for example, a protein network for a specific tissue of interest.

The stringApp is thus also useful beyond analysis of user-provided high-throughput data. For example, one can easily perform a disease query to retrieve a network of proteins known to be involved in a given disease, use network expansion to obtain novel candidates, filter them by expression in disease-relevant tissues, and highlight the druggable targets from Pharos.

### Automation of analyses

In addition to having a graphical user interface, which we have focused on in this paper, the stringApp also supports the Cytoscape Automation feature, which allows scripted execution of STRING analyses within Cytoscape. This command interface can be used in a variety of ways. First, it is accessible through the Command Tool, which provides an interactive command line as well as the ability to execute Cytoscape script files. Second, the commands can be used from web pages viewed in the built-in Cytoscape web browser, as illustrated in the online stringApp training material (https://jensenlab.org/training/stringapp/). Third, the cyREST app^23^ enables other programs to control Cytoscape through an API, which in turn allows stringApp analyses to be scripted from R using the BioConductor package RCy3^24^ or from Python using package py2cytoscape^23^. A tutorial on the latter can be found in the Cytoscape Automation training material (https://git.io/RstringAppTutorial).

### Open challenges in network visualization of proteomics data

There are still several open challenges in network visualization of MS-based proteomics data, which are in no way specific to stringApp but also not addressed by it.

The proteolytic cleavage of the proteins, typically with trypsin, results in peptides that do not map uniquely to specific proteins. Instead, these peptides are generally mapped to so-called protein groups, which consist of multiple proteins from which the peptides could have originated. How to best represent this ambiguity in a protein network is not clear; options include choosing a single representative protein for each group, showing all proteins from each group, or constructing network nodes that fuse all interaction evidence for the proteins in a protein group.

Another challenge relates specifically to data on post-translational modifications. Since each protein can have multiple post-translational modifications on different sites, an MS dataset may show that some sites on a protein are up-regulated while others are down-regulated. Since a protein network will have only a single node for each protein, visualization of site-specific data requires multiple values to be shown on each node, for example, in the form of a donut plot. However, this visualization will result in information overload if used directly on large networks. Visualization of networks with complex data overlays, such as time courses or site-specific data, might be achieved by separating the data from the network view and using interactive techniques to identify sub-networks of interest^25^.

## Supporting Information

Figure of the clustered protein association network of proteins with significantly regulated phosphorylation sites (Supporting Information 1, Figure S1, PDF)

Table of the significant phosphorylated sites identified by Francavilla *et al.*^13^ (Supporting Information 2, Table S1, XLSX)

## Acknowledgments

The authors thank Helen V. Cook for testing the software and giving detailed feedback on the user interface. The authors also thank Damian Szklarczyk, Michael Kuhn, and Christian von Mering for implementing the changes to the STRING and STITCH APIs, which were needed to support stringApp.

This work was funded by the Novo Nordisk Foundation (NNF14CC0001), the Danish Council for Independent Research (DFF-4005-00443), and the National Institutes of Health (NIH) Illuminating the Druggable Genome Knowledge Management Center (U54 CA189205 and U24 224370), NIH NIGMS P41 GM103504, and grant number 2018-183120 from the Chan Zuckerberg Initiative DAF, and advised fund of the Silicon Valley Community Foundation.

## References

(1) Szklarczyk, D.; Morris, J. H.; Cook, H.; Kuhn, M.; Wyder, S.; Simonovic, M.; Santos, A.; Doncheva, N. T.; Roth, A.; Bork, P.; et al. The STRING Database in 2017: Quality-Controlled Protein-Protein Association Networks, Made Broadly Accessible. Nucleic Acids Res. 2017, 45 (D1), D362–D368.

(2) Shannon, P.; Markiel, A.; Ozier, O.; Baliga, N. S.; Wang, J. T.; Ramage, D.; Amin, N.; Schwikowski, B.; Ideker, T. Cytoscape: A Software Environment for Integrated Models of Biomolecular Interaction Networks. Genome Res. 2003, 13 (11), 2498–2504.

(3) Cline, M. S.; Smoot, M.; Cerami, E.; Kuchinsky, A.; Landys, N.; Workman, C.; Christmas, R.; Avila-Campilo, I.; Creech, M.; Gross, B.; et al. Integration of Biological Networks and Gene Expression Data Using Cytoscape. Nat. Protoc. 2007, 2 (10), 2366–2382.

(4) Morris, J. H.; Apeltsin, L.; Newman, A. M.; Baumbach, J.; Wittkop, T.; Su, G.; Bader, G. D.; Ferrin, T. E. clusterMaker: A Multi-Algorithm Clustering Plugin for Cytoscape. BMC Bioinformatics 2011, 12, 436.

(5) Tay, A. P.; Pang, C. N. I.; Winter, D. L.; Wilkins, M. R. PTMOracle: A Cytoscape App for Covisualizing and Coanalyzing Post-Translational Modifications in Protein Interaction Networks. J. Proteome Res. 2017, 16 (5), 1988–2003.

(6) Pletscher-Frankild, S.; Palleja, A.; Tsafou, K.; Binder, J. X.; Jensen, L. J. DISEASES: Text Mining and Data Integration of Disease-Gene Associations. Methods San Diego Calif 2015, 74, 83–89.

(7) Szklarczyk, D.; Santos, A.; von Mering, C.; Jensen, L. J.; Bork, P.; Kuhn, M. STITCH 5: Augmenting Protein-Chemical Interaction Networks with Tissue and Affinity Data. Nucleic Acids Res. 2016, 44 (D1), D380–D384.

(8) Binder, J. X.; Pletscher-Frankild, S.; Tsafou, K.; Stolte, C.; O’Donoghue, S. I.; Schneider, R.; Jensen, L. J. COMPARTMENTS: Unification and Visualization of Protein Subcellular Localization Evidence. Database J. Biol. Databases Curation 2014, 2014, bau012.

(9) Palasca, O.; Santos, A.; Stolte, C.; Gorodkin, J.; Jensen, L. J. TISSUES 2.0: An Integrative Web Resource on Mammalian Tissue Expression. Database J. Biol. Databases Curation 2018, 2018, bay003.

(10) Nguyen, D.-T.; Mathias, S.; Bologa, C.; Brunak, S.; Fernandez, N.; Gaulton, A.; Hersey, A.; Holmes, J.; Jensen, L. J.; Karlsson, A.; et al. Pharos: Collating Protein Information to Shed Light on the Druggable Genome. Nucleic Acids Res. 2017, 45 (D1), D995–D1002.

(11) Fielding, R. T. Chapter 5: Representational State Transfer (REST). In Architectural Styles and the Design of Network-based Software Architectures; University of California, Irvine, 2000; pp 76–106.

(12) Hobohm, U.; Scharf, M.; Schneider, R.; Sander, C. Selection of Representative Protein Data Sets. Protein Sci. Publ. Protein Soc. 1992, 1 (3), 409–417.

(13) Francavilla, C.; Lupia, M.; Tsafou, K.; Villa, A.; Kowalczyk, K.; Rakownikow Jersie-Christensen, R.; Bertalot, G.; Confalonieri, S.; Brunak, S.; Jensen, L. J.; et al. Phosphoproteomics of Primary Cells Reveals Druggable Kinase Signatures in Ovarian Cancer. Cell Rep. 2017, 18 (13), 3242–3256.

(14) Enright, A. J.; Van Dongen, S.; Ouzounis, C. A. An Efficient Algorithm for Large Scale Detection of Protein Families. Nucleic Acids Res. 2002, 30 (7), 1575–1584.

(15) Morris, J. H.; Kuchinsky, A.; Ferrin, T. E.; Pico, A. R. enhancedGraphics: A Cytoscape App for Enhanced Node Graphics. F1000Research 2014, 3, 147.

(16) The UniProt Consortium. UniProt: The Universal Protein Knowledgebase. Nucleic Acids Res. 2017, 45 (D1), D158–D169.

(17) Stelzer, G.; Rosen, N.; Plaschkes, I.; Zimmerman, S.; Twik, M.; Fishilevich, S.; Stein, T. I.; Nudel, R.; Lieder, I.; Mazor, Y.; et al. The GeneCards Suite: From Gene Data Mining to Disease Genome Sequence Analyses. Curr. Protoc. Bioinforma. 2016, 54, 1.30.1–1.30.33.

(18) Harrower, M.; Brewer, C. A. ColorBrewer.org: An Online Tool for Selecting Colour Schemes for Maps. Cartogr. J. 2003, 40 (1), 27–37.

(19) McGrath, C.; Krackhardt, D. Visualizing Complexity in Networks : Seeing Both the Forest and the Trees; 2003.

(20) Muñoz, M. J.; de la Mata, M.; Kornblihtt, A. R. The Carboxy Terminal Domain of RNA Polymerase II and Alternative Splicing. Trends Biochem. Sci. 2010, 35 (9), 497–504.

(21) Brown, L. A.; Williams, J.; Taylor, L.; Thomson, R. J.; Nolan, P. M.; Foster, R. G.; Peirson, S. N. Meta-Analysis of Transcriptomic Datasets Identifies Genes Enriched in the Mammalian Circadian Pacemaker. Nucleic Acids Res. 2017, 45 (17), 9860–9873.

(22) Huang, J.; Vendramin, S.; Shi, L.; McGinnis, K. M. Construction and Optimization of a Large Gene Coexpression Network in Maize Using RNA-Seq Data. Plant Physiol. 2017, 175 (1), 568–583.

(23) Ono, K.; Muetze, T.; Kolishovski, G.; Shannon, P.; Demchak, B. CyREST: Turbocharging Cytoscape Access for External Tools via a RESTful API. F1000Research 2015, 4, 478.

(24) Shannon, P. T.; Grimes, M.; Kutlu, B.; Bot, J. J.; Galas, D. J. RCytoscape: Tools for Exploratory Network Analysis. BMC Bioinformatics 2013, 14, 217.

(25) Hochheiser, H.; Shneiderman, B. A Dynamic Query Interface for Finding Patterns in Time Series Data. In CHI ‘02 extended abstracts on Human factors in computing systems - CHI ‘02; ACM Press: Minneapolis, Minnesota, USA, 2002; p 522002E

